# A human kinase yeast array for the identification of kinases modulating phosphorylation-dependent protein-protein interactions

**DOI:** 10.1101/2021.11.18.469002

**Authors:** Stefanie Jehle, Natalia Kunowska, Nouhad Benlasfer, Jonathan Woodsmith, Gert Weber, Markus C. Wahl, Ulrich Stelzl

## Abstract

Protein kinases play an important role in cellular signaling pathways and their dysregulation leads to multiple diseases, making kinases prime drug targets. While more than 500 human protein kinases are known to collectively mediate phosphorylation of over 290,000 S/T/Y sites, the activities have been characterized only for a minor, intensively studied subset. To systematically address this discrepancy, we developed a human kinase array in *Saccharomyces cerevisiae* as a simple readout tool to systematically assess kinase activities. For this array, we expressed 266 human kinases in four different *Saccharomyces cerevisiae* strains and profiled ectopic growth as a proxy for kinase activity across 33 conditions. More than half of the kinases showed an activity-dependent phenotype across many conditions and in more than one strain. We then employed the kinase array to identify the kinase(s) that can modulate protein-protein-interactions (PPIs). Two characterized, phosphorylation-dependent PPIs with unknown kinase-substrate relationships were analyzed in a phospho-yeast two-hybrid assay. CK2α1 and SGK2 kinases can abrogate the interaction between the spliceosomal proteins AAR2 and PRPF8 and NEK6 kinase was found to mediate the estrogen receptor (ERα) interaction with 14-3-3 proteins. The human kinase yeast array can thus be used for a variety of kinase activity-dependent readouts.

## Introduction

Protein kinases represent one of the largest protein families in eukaryotes, with the 518 annotated human kinases comprising about 2% of the protein coding genome [1,2]. The kinase domain catalyzes the phosphorylation of mainly serine, threonine and tyrosine residues. Already, over 290,000 phosphorylation sites in human proteins have been reported (phosphosite.org; 09/21). Phosphorylation of a protein can have a wide range of molecular effects, such as changes in activation status, conformation, localization, or interaction patterns. Phosphorylation underpins cellular signaling pathways and regulates diverse processes such as transcription, metabolism, the cell cycle, and proliferation. Consequently, dysregulated protein kinases often drive disease phenotypes, and elucidating kinase activity profiles [3], their substrate relationships [4–6] and their effects on cellular processes and phenotypes [7] is imperative to better understand the mechanisms of disease progression. Multiple kinase inhibitors are used as first line therapy in many cancers as well as in other diseases [8]. Despite the large number of human kinases with a pivotal role in cellular signaling, there is a well-documented research bias towards a small set of highly-studied kinases such as p38a, SRC, EGFR, PKA, ERK1, KDR, ABL, CDK1 or KIT [9,10]. Studying kinase activity within cells is difficult, as kinases exhibit overlapping substrate specificity and differential expression, depending on the cell type, cell cycle phase, or subcellular localization, and have order(s) of magnitude differences in enzymatic activities. Furthermore, kinases form complex signaling networks that contain redundancy and feedback loops. Therefore, assaying kinase activity is typically done with purified kinase domains *in vitro* or with engineered biosensors [11–14]. However, a versatile simple readout to analyze wild type kinase activities in parallel at the kinome scale would be a valuable complement to current approaches.

Human kinases have been shown to be active in yeast. For example, expression of human serine/threonine (S/T) cell cycle kinase CDK1 suppresses lethality of *cdc28* kinase mutant alleles in yeast [15,16]. The yeast *kin28* deletion strain could be complemented by seven human paralog kinases [17]. Also active tyrosine (Y) kinases have been expressed in yeast to reveal kinase regulation mechanisms [18], identify protein-protein interactions [19] and study mechanism of kinase specificity [5]. Elevated expression of active human kinases quickly causes yeast growth defects. When expressing v-SRC, yeast growth was diminished [20–23] and later reports confirmed that human protein tyrosine kinase toxicity in yeast can be explained by aberrant phosphorylation of yeast proteins [24–26]. Likewise, when human S/T kinase AKT1 is expressed in yeast it is phosphorylated on T308 and S473 and thus activated [27]. When AKT1 activity is increased either through oncogenic mutations or co-expression of the catalytic subunit of PI3 kinase, it causes increased growth inhibition of yeast cells [28]. In order to set up a drug screening platform in yeast, Sekigawa et al. utilized a human cDNA library which also contained human kinases for overexpression screening. About 23% of the tested kinases, including serine/threonine kinases, caused a visible growth defect of yeast under standard growth conditions [29]. Recently, screening of >500 human kinases revealed 28 kinases with strong overexpression-associated toxicity in yeast, that was further leveraged to screen for genetic suppressors of this growth defect [30].

Here, we aimed to systematically exploit the yeast growth phenotype caused by overexpression of active human kinases as an activity readout. While 15% of the 6100 *S. cerevisiae* genes are essential under standard growth conditions, approximately another 63% of the *S. cerevisiae* open reading frames (ORFs) were required for full growth when tested under more than 1000 different conditions [31]. In addition, essentiality of *S. cerevisiae* genes is strongly dependent on genetic interactions, which vary within different genotype of the strains [32]. This can be explained by the cellular protein interaction networks underlying the genotype to phenotype relationship, which are strongly modulated through genetic and environmental variation [33,34]. Therefore, we hypothesized that the deleterious effects of aberrant phosphorylation of yeast proteins by human kinases may be further enhanced under non-standard culture conditions and in different genomic backgrounds of different *S. cerevisiae* yeast strains.

We developed a human protein kinase yeast array that comprises more than 50% of the annotated human kinome and utilizes ectopic yeast growth as a simple readout for kinase activity in a living organism. Yeast growth is assayed with four strains with different genomic backgrounds and across a large variety of environmental conditions. Kinase activity is demonstrated growth assays in comparison with kinase-dead mutant versions and by assaying phosphorylation of yeast proteins directly with different phospho-substrate antibodies on whole yeast protein lysates. The yeast array is a versatile tool to broadly capturing human kinase activity which can be coupled to any yeast assay available. Here, we demonstrate its utility by applying the array in identification of the kinases modulating known phosphorylation dependent PPIs.

## Results

### An array of human protein kinases in budding yeast

To construct a large human kinase array in yeast and leverage yeast growth phenotypes as kinase activity readout under various conditions and different genetic backgrounds, we collected ORFs for 266 human protein kinases (**Figure 1A**). As a control group for the kinases, we employed 80 proteins, including 10 kinase-dead versions and 70 unrelated proteins without kinase domains. 92% of the proteins in the kinase set and control set were covered with two or more ORFs. We subcloned the ORFs into two yeast expression vectors, both under a weak Cu^2+^ inducible yeast promoter, either with or without a nuclear localization sequence [5,19], arriving at a total of 1017 different plasmids. The 266 protein kinases represented over a half of the known human kinome from all major groups and families of the Human Kinome Tree [2] (**Figure 1B**).

**Figure 1:**
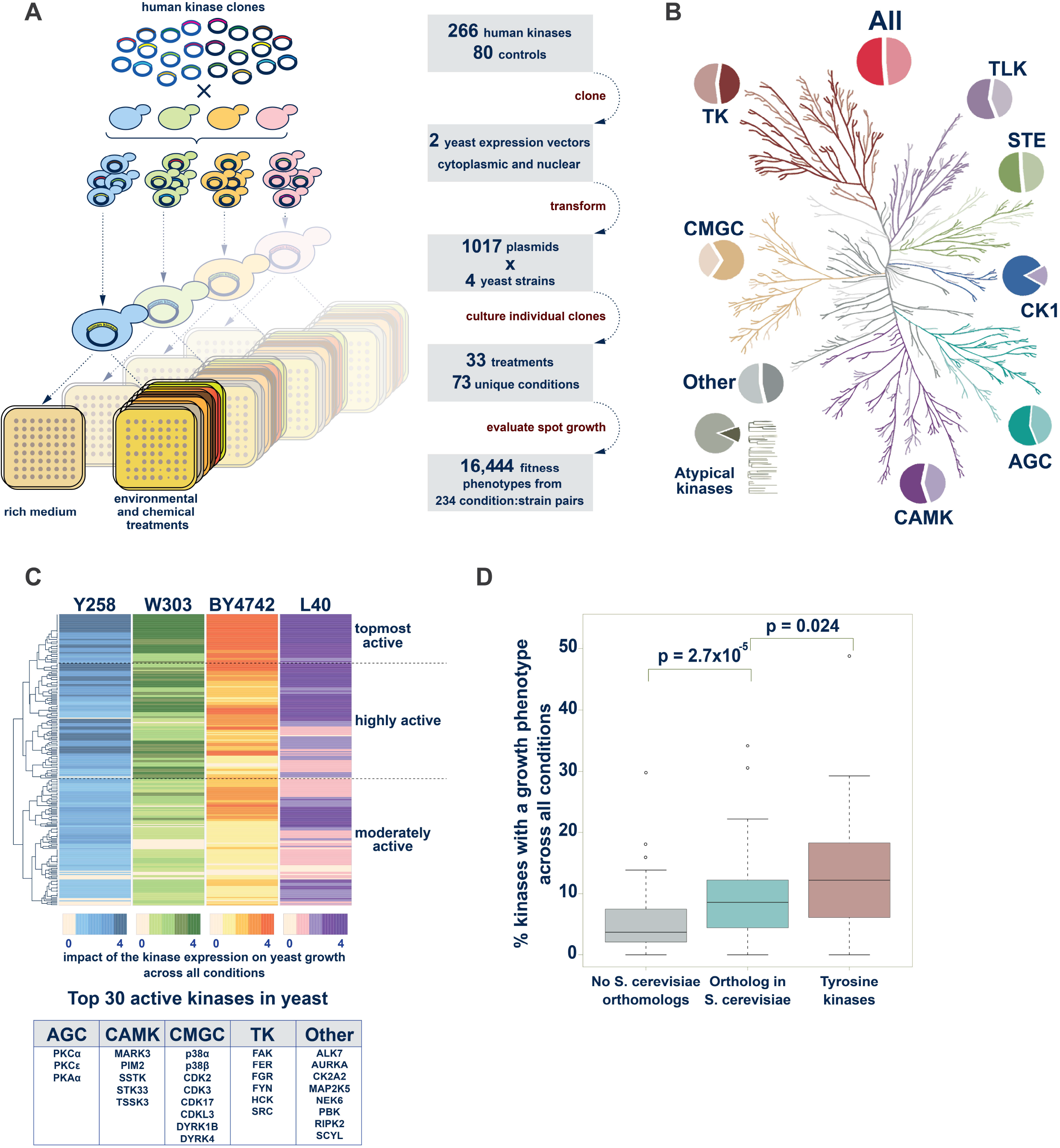
The human kinase yeast array. **A.** Schematic representation of the principle of the kinase yeast array and a flow diagram showing the construction of the human kinase array. **B.** The Human Kinome Tree representing 518 kinases in nine families (modified from [2] using CORAL at http://phanstiel-lab.med.unc.edu/CORAL/). Pie charts indicate the fraction of kinases contained in the human kinase yeast array. In total, 266 human protein kinases and 52% of the human kinome is represented in the array. **C.** Hierarchical clustering of the results of the yeast growth phenotype screen. In each strain, the 266 kinases were binned (0-4) according to the number of conditions in which the kinase caused a growth phenotype. Hierarchical clustering revealed sets of topmost (52), highly (98) and moderately active (116) kinases. Top 30 active kinases causing many growth phenotypes over all conditions-strain pairs are listed. **D.** Comparison of the fraction of kinases that result a growth phenotype over all conditions-strain pairs. Kinases are grouped into 179 (67%) with a yeast ortholog and 87 (33%) human protein kinases without orthologous yeast kinases. The latter group includes 41 tyrosine kinases and 46 S/T kinases which were separated in the analysis. The former group contains both one-to-many and many-to-many orthologous kinases. Tyrosine kinases strongly inhibited yeast growth and human kinases that are orthologous to yeast kinases were more likely to perturb yeast growth than human kinases without a yeast ortholog.

The basic underlying principle of our kinase yeast array approach is that kinase activity will lead to aberrant phosphorylation of yeast proteins and thereby impair the yeast growth. In an initial test, the induction of kinase expression at 20 μM Cu^2+^ resulted a yeast growth phenotype for 18% of the kinases. To be able to assay kinase activity using growth phenotypes as broadly as possible, we used four *S. cerevisiae* strains with different genomic backgrounds exhibiting different protein expression and growth phenotypes: the commonly used haploid laboratory strain BY4742 (MATα) [35,36], the haploid yeast two-hybrid (Y2H) strain L40c (MATa) [37,38], the diploid strain W303 (MATa/MATα) [39], and the haploid protein expression strain Y258 (MATa) [40]. They were individually transformed with the two sets of kinase plasmids. We then cultured these strains in parallel in 384-array format on agar under various growth conditions (**Figure 1A**). The conditions included carbon sources other than glucose, the presence of detergents or denaturing agents, environmental triggers (pH, temperature), and addition of metal ions, osmotic stress agents, and other chemicals (such as MnCl_2_, K_3_Fe(CN)_6_, or LiCl) to the growth media. Together, 33 unique treatments were assayed with additives in various concentrations adding up to a total of 73 different conditions in which at least one of the four strains were evaluated successfully (**Supplementary Table 1**). The evaluation of over 300,000 yeast spots for 266 kinases revealed that many kinases show growth reduction in at least one condition for one strain. Growth was evaluated on a 0-3 scale, reflecting none, weak, moderate, to severe growth reduction, respectively. Conditions which reduced growth of more than 175 (17%) colonies were excluded from further analysis. A total of 243 condition-strain pairs were considered, with 41 conditions evaluated for all four strains and a total of 70, 62, 64, and 47 conditions for BY4742, L40c, W303, and for Y258, respectively **(Supplementary Table 2)**. The use of different lab strains further increased sensitivity. In total 25, 36, 49, and 89 kinases with severe growth reduction were detected in at least 4, 3, 2 or 1 yeast strain respectively.

The 266 human protein kinases were then clustered based on their activity (**Figure 1C**). For that, the kinases were grouped in four bins, according to number of conditions where they exhibit a growth phenotype and ranked separately for each strain. Hierarchical clustering revealed sets of kinases that were active in many conditions in one strain, two, or more strains, respectively. We grouped kinases according to their activity profiles across the four strains into three activity groups, from topmost active through highly to moderately active kinases (**Figure 1C, Supplementary Table 2**). In agreement with previous observations [5], the topmost active kinases included cytoplasmic tyrosine kinases, which caused strong growth reduction when overexpressed in yeast. They also included two PKC family members (alpha and epsilon), two p38 MAP kinase family members (alpha and beta) as well as the cell cycle kinases CDK2, CDK3. Likewise, MARK3, RIPK2, NEK6, CDK17, CSNK2A2, AURKA, DYRK4, DYRK1B, and PKA were members of the high activity group, with their expression frequently resulting in growth reduction (**Figure 1C**, box with the top 30).

Next, we compared the percentage of strong growth phenotypes caused by human kinases which have homologous kinase(s) in *S. cerevisiae* with human kinases that do not have homologs in yeast. Tyrosine kinases were treated as a separate group because tyrosine kinase signaling is a hallmark of multicellularity and did not evolve in yeast. Aberrant tyrosine phosphorylation can be toxic in yeast and it was suggested that organisms with evolved tyrosine signaling have a substantially reduced tyrosine content to prevent adverse tyrosine phosphorylation [41]. The results illustrate that the percentage of kinases causing a strong growth phenotype per condition is highest amongst the tyrosine kinases. The percentage of serine/threonine kinases causing a strong growth phenotype per condition is higher for the human kinases with a homolog in yeast than for those without (**Figure 1D**). This on one hand confirms that high level tyrosine phosphorylation in yeast is harmful, and on the other hand showed a significant trend suggesting that conserved S/T-kinases are more likely to interfere with essential yeast cellular process perturbing yeast growth than heterologous expression of non-conserved kinases.

### Kinase activity is critical for yeast growth phenotypes

Evaluation of the yeast growth phenotype under a variety of growth conditions allowed monitoring the effects of the expression of individual human kinases through a simple readout such as colony size on agar. However, heterologous expression of any protein in yeast as such can affect yeast growth, independently of the activity of the expressed protein. Therefore, we asked whether kinase phosphorylation activity and not human protein overexpression per se drives our phenotypic readout. We first compared the number of growth phenotypes caused by the human protein kinases with the number of growth phenotypes observed with the control set of proteins without kinase activity. The statistical comparison of the number of observed phenotypes for the two groups reveals that kinase expression resulted in growth defects much more often than expression of unrelated proteins or phosphorylation-inactive kinase versions (**Figure 2A**).

**Figure 2:**
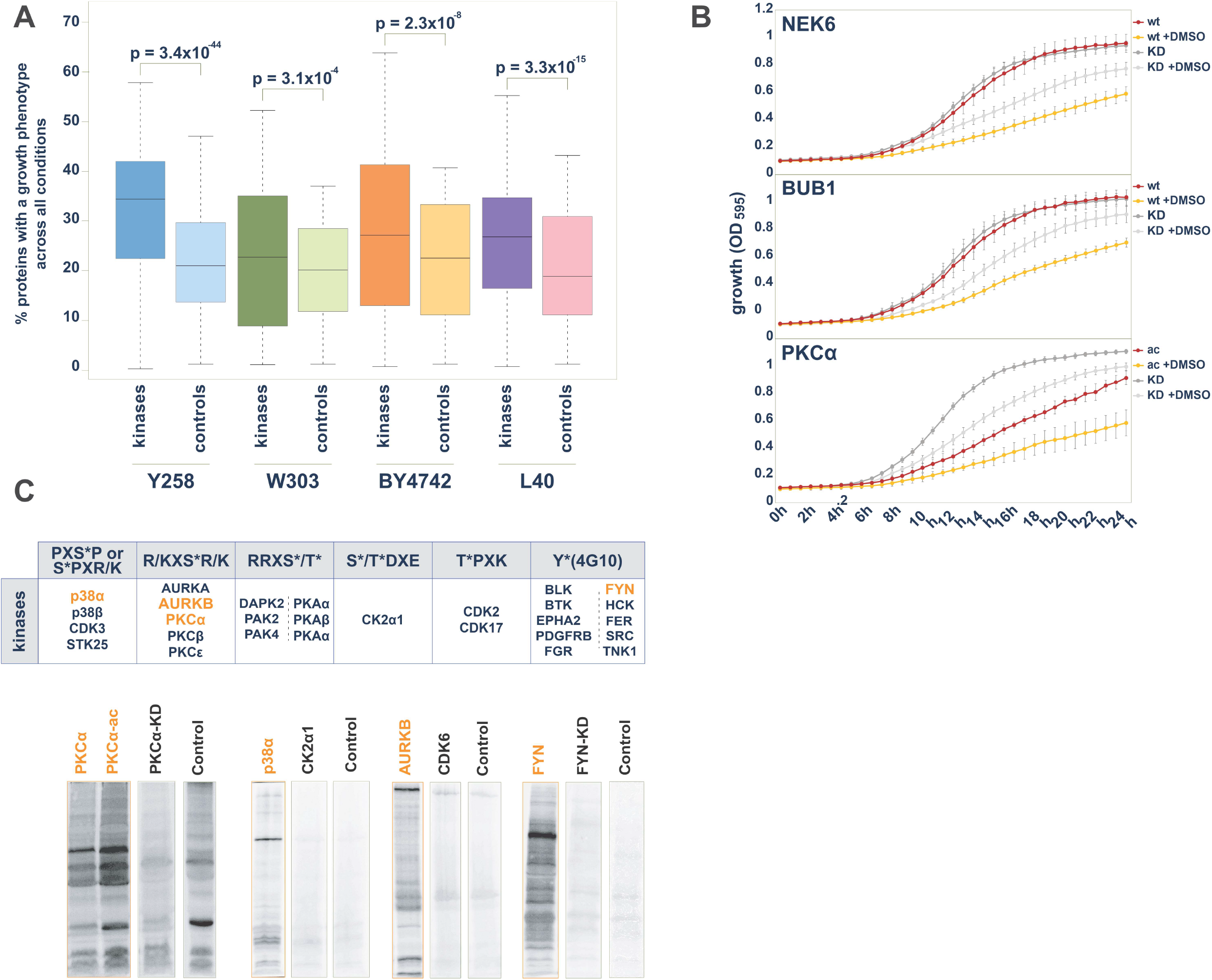
The yeast growth phenotype is a readout for the kinase phosphotransferase activity. **A.** Statistical comparison of the number of yeast growth phenotypes between kinase group and control protein group. In each strain, across all conditions, the percentage of kinases that cause a phenotype upon exogenous expression is significantly higher than the percentage of control proteins, than cause a growth phenotype upon overexpression. **B.** Growth comparisons of selected kinases with the kinase-dead mutant match. 24h growth curves (OD_595_) from liquid culture of yeast strains (W303) expressing human kinase in the presence or absence of 3.2% (v/v) DMSO. Upper panel: BUB1: BUB1 wild type, BUB1-KD: NEK6(K821M); middle panel: NEK6: NEK6 wild type, NEK6-KD: NEK6(K74M&K75M); lower Panel: PKCα-ac: PKCα(A25E), PKCα-KD: PKCα(K368R) **C.** Human protein kinases phosphorylate large sets of yeast proteins. Upper panel: In total, 27 human protein kinases showed activity towards yeast proteins visible on western blots through enhanced reactivity with phospho-motif antibodies on yeast proteins in comparison to other human kinases, kinase-dead mutant versions or vector control samples. Lower panel: Examples of western blots with equal amount of whole yeast cell lysate from strains expressing human kinases loaded. When developed with the indicated phospho-motif recognizing antibodies, immunoreactive bands indicated phosphorylation of yeast proteins.

To experimentally validate the impact of kinase activity on observed growth phenotypes in the yeast array, we monitored growth measuring the optical density of liquid cultures of wild type kinase in parallel with kinase-dead mutant version over 24 hours. The results for BUB1 and NEK6 illustrate that there is no difference in yeast growth under standard growth condition in this assay. The addition of 3.2% DMSO to the media inhibited yeast growth, observable through a shallower growth curve and lower maximum cell density. In the presence of DMSO, the expression of the wildtype kinase of BUB1 or NEK6 diminishes yeast growth in comparison to the respective BUB1 (K821M) and NEK6 (K74M&K75M) kinase-dead mutants (**Figure 2B**). The comparison of yeast growth upon expressing either the activated form of PKCα (A25E) or the PKCα (K368R) kinase-dead mutant revealed pronounced differences under both standard and 3.2% DMSO growth conditions (**Figure 2B**). These results of this liquid growth assay were in good agreement with the results of the solid media screen, confirming that the kinase activity is required to inhibit yeast growth.

To directly show kinase activity in the yeast strains we assayed kinases for their ability to phosphorylate endogenous yeast proteins using phospho-specific antibodies. To this end, whole cell lysates from yeast expressing human kinases were subjected to SDS-PAGE and western blotting and probed with either a general phospho-tyrosine recognizing antibody (4G10) or five different phospho-substrate antibodies which recognize phosphorylated S/T-sites preferentially in a certain amino acids sequence context (**Figure 2C)**. Equal amount of whole cell lysate was probed comparing different kinases as well as kinases with the respective kinase-dead mutant variants. As illustrated for PKCα, p38α, AURKB, and FYN in **Figure 2C**, diverse patterns of immunoreactivity were obtained when probed with indicated antibodies. Most tyrosine kinases phosphorylated yeast proteins, albeit with different sensitivity and specificity [5]. Endogenous serine and threonine phosphorylation in yeast is high. Therefore, phosphorylation patterns were distinguishable from background signals for only a subset of the tested kinases and antibodies. In total, 27 human protein kinases showed phosphorylation activity towards yeast proteins with one of the six available antibodies (**Figure 2C**). These experiments showed that human kinases are active in yeast and phosphorylate yeast proteins.

### The kinase array is a versatile tool to study human kinase activities

Our array set up to study kinase activities revealed a set of 150 highly active kinases (topmost and highly active). This subset of the human kinome included representatives from all major kinase families (**Figure 3A**). Many well characterized kinases are known cancer drivers, and more than 340 human kinases are associated with at least one type of cancer [42]. 95 of the highly active kinases overlap with this cancer kinase set (**Figure 3B)**. One major strength of the yeast kinase array is that it also covers a large number of less well studied or difficult to study kinases. 162 kinases, nearly one third of the kinome, have been designated by the NIH as being poorly understood. As their role or biological function is largely unknown they are termed ‘dark kinases’ [10]. Our array contained 72 dark kinases, 35 of which were in the group of highly active kinases (**Figure 3C**).

**Figure 3:**
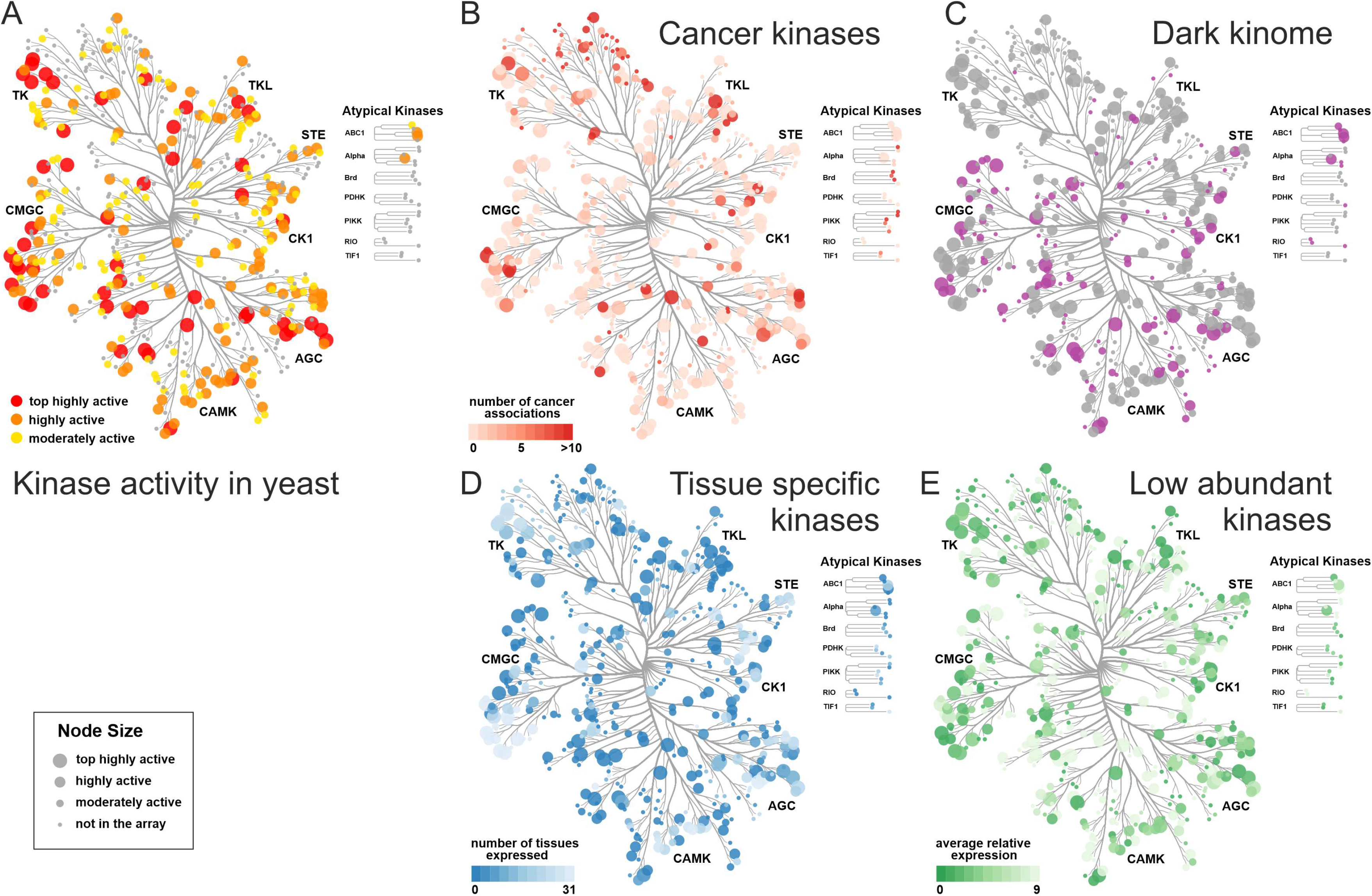
Representation of active human kinases in the yeast array. **A.** Kinase activity in yeast. Kinase activity is played on the human kinome tree showing that all kinase families contain highly active kinases members. **B.** Cancer kinases. The color-coded annotated cancer kinases overlap with of highly active kinases (larger node size) in the array. 168 kinases in the array are linked to at least one cancer type [42]. **C.** Dark kinases. Dark kinases [10] in the kinome are color coded in purple, the tree shows the overlap of 35 highly active kinases (larger node size) and the total of 72 dark kinases represented in the array. **D.** Tissue specific kinases. Normalized proteomics data [43] from 31 tissues show that some kinases are only expressed in a few tissues. Many tissue specific kinases (darker color) belong to the highly active kinases (larger node size) in the yeast array. **D**. Low abundant kinases. Average relative expression levels over 31 tissues from PaxDB are color coded. Many kinases with low expression in mammalian cells (darker color) belong to the highly active kinases (larger node size) in the yeast array. Figures **A-E** were prepared using CORAL at http://phanstiel-lab.med.unc.edu/CORAL/.

In general, studying kinases in their cellular environment is challenging, as kinase perturbation results in widespread pleiotropic cellular effects. In addition, many kinases are also highly tissue specific and/or low abundant, posing further challenges for the characterization of activities. We used protein abundance data from 31 tissues from the PAXdb (Protein Abundance Database), that provides normalized relative protein abundance data across many diverse mass spectrometry-based proteome studies [43]. The kinome tree view shows that tissue specific and low-abundance kinases are well covered in our array (**Figure 3D**, tissue expression, **Figure 3E**, average protein abundance). For example, two members each of the TSSK, MARK or DRYK kinases, all highly active in yeast, belong to the dark, tissue-specific group of kinases with low expression. The activity of human kinases in the yeast array is largely independent of endogenous expression levels in human cells, thus enabling to study all candidate kinases equally well.

### Phospho-yeast two-hybrid for identification kinases modulating PPIs

In our system, the activities of many human kinases can be monitored by growth reduction under varying conditions. However, low to moderate expression of human kinases under standard conditions does not impair yeast growth. Therefore, this kinase expression array opens avenues to alternative screening approaches, where the activity of a kinase can be assessed when coupled to any functional assay or readout in yeast. Cellular assays can involve drug inhibition/resistance selection, expression of a reporter such as luciferase, or expression of a fluorescent or epitope-tagged protein, or functional genomics readouts such sequencing or mass spectrometry [44]. To explore the screening possibilities offered by the kinase array, we applied a phospho-yeast two-hybrid system [19] to identify kinases that modulate phosphorylation-dependent protein-protein interactions. The phosphorylation of one protein can either abolish its ability to interact with its partner, resulting in a loss of interaction, or it can enable the interaction with a second protein leading to a gain of interaction (**Figure 4A**). Both types of phosphorylation-dependent PPI switches require an active kinase facilitating the phosphorylation of one of the interaction partners. In the first setup where phosphorylation abolishes the Y2H interaction, active kinases are expressed from a third plasmid and phosphorylate either the bait or prey protein, resulting in reduced yeast growth on selective media. Conversely, the kinase activity can also promote yeast growth on selective media when the kinase activity enables a protein complex formation (**Figure 4A**). Modulation of protein interactions by phosphorylation has been widely described in the literature [45], however it often remains unclear which kinase(s) are responsible for interaction regulation. There are many “kinase-orphan” phosphorylation-dependent interactions described in the literature [46] but no systematic approach tailored towards identifying kinases directly involved in phospho-mediated modulation of protein interactions has been reported. Our kinase array allows us to identify such kinases with a phospho-Y2H matrix approach (**Figure 4B**). A bait-prey pair of human proteins is co-expressed in one yeast strain (L40c **matα**) and mated with each of the L40c **mat***a* strains in the kinase array. The active kinases modulating the interaction of the two proteins of interest are then identified by altered yeast growth patterns (**Figure 4A**). To test the utility of this array in identifying biologically relevant kinase activities, we screened two well characterized phosphorylation-dependent interactions, which represent the two cases and for which the responsible modulatory kinases are as yet unknown.

**Figure 4:**
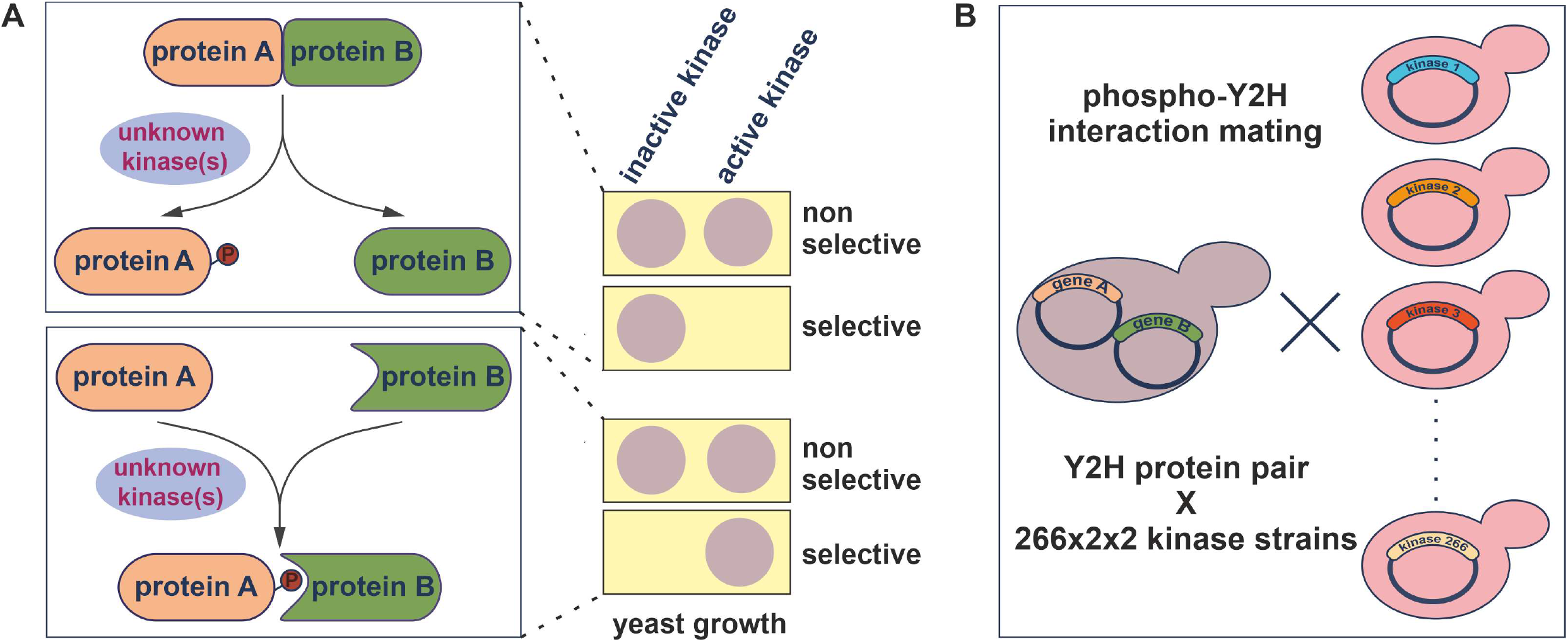
The human kinase yeast array as a tool to identify the kinase(s) modulating a phosphorylation-dependent PPIs. **A.** Two distinct cases of phosphorylation-dependent interaction modulation. Loss of interaction: the phosphorylation of one protein (A) by the active kinase results in the loss of interaction with the second protein (B) and therefore in no growth on selective media. Gain of interaction: Co-expression of an active kinase with a Y2H bait-prey protein pair can lead to phosphorylation of one protein partner (A) promoting the pS/T-dependent interaction with protein (B). In the phospho-Y2H approach, gain of interaction results in yeast growth on selective media. **B.** Systematic phospho-Y2H interaction mating approach to identify the active kinase(s) modulating phosphorylation-dependent PPI. One yeast strain (L40matα) co-expressing the interacting bait and prey proteins and was mated against the 266 kinases in the array (L40mata) in a pairwise manner. The growth patterns of the diploid yeast strains that express bait, prey and one kinase from a third plasmid, indicate phospho-modulation of protein interactions.

### Kinase-dependent inhibition of the spliceosomal AAR2 - PRPF8 interaction

PRP8 is a major constituent of the spliceosome, delivered as part of the U5 snRNP complex during the spliceosomal assembly cycle. AAR2 is a U5 snRNP and U4/U6-U5 tri-snRNP assembly factor. In yeast, AAR2 is thought to dissociate from PRP8 at late assembly stages based on a phosphorylation trigger freeing PRP8 to allow for BRR2 binding (**Figure 5A**). While these events, including the crucial AAR2 phosphorylation site at serine 253, are well characterized in yeast [47–49], the kinase(s) responsible for the phosphorylation are unknown.

**Figure 5:**
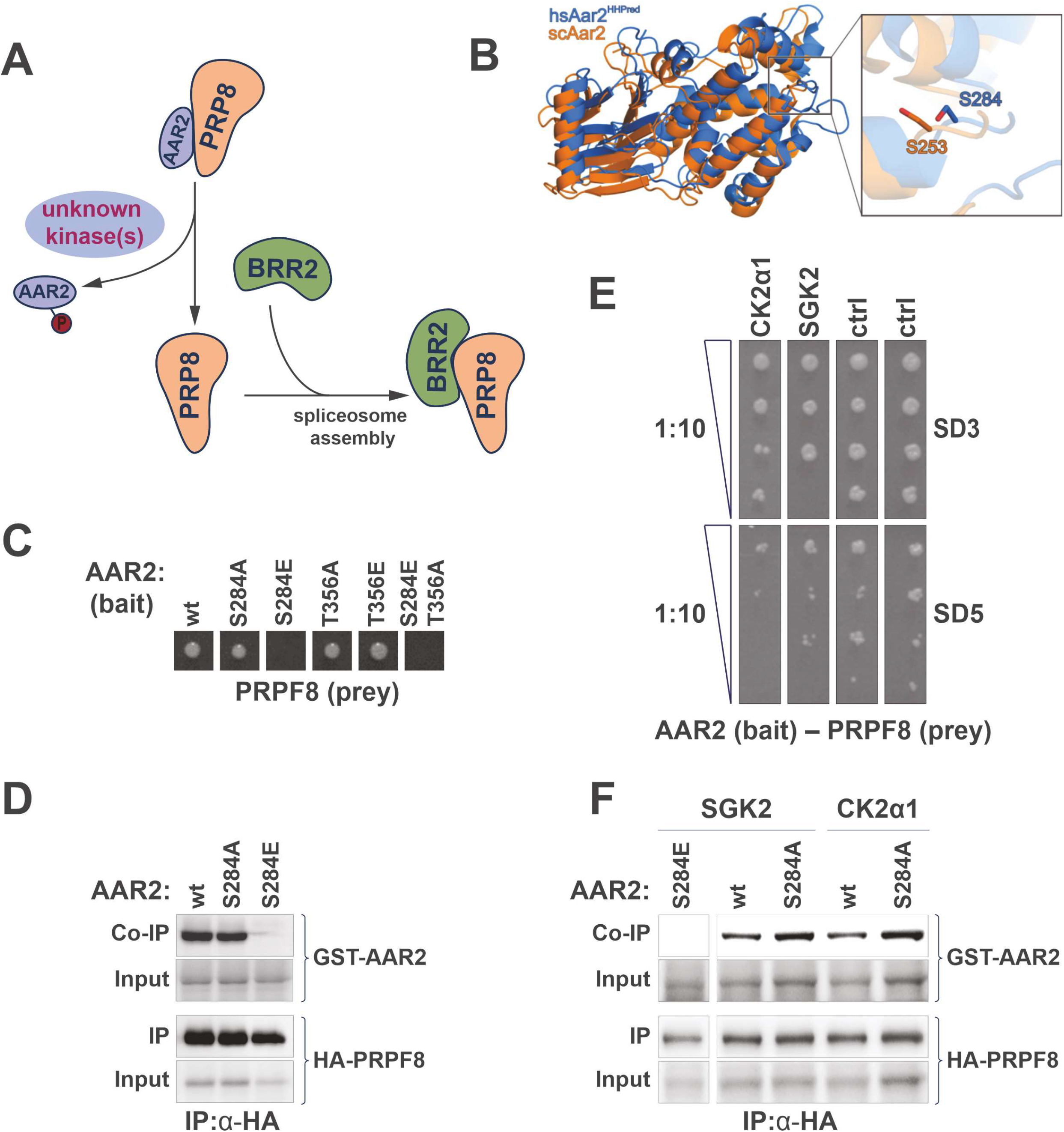
Identification of human S/T kinases modulating the AAR2-PRP8 protein interaction. **A.** Schematic of phospho-dependent interaction of the spliceosomal AAR2 and PRP8. AAR2, a spliceosomal assembly protein, and PRP8 from a stable complex, that dissociates upon phosphorylation of AAR2. PPR8 is then free to bind the spliceosomal interaction partner protein BRR2 [48,49]. This regulatory interaction switch is thought to be conserved between yeast and human. A protein kinase that can modulate the dissociation of AAR2 from PRP8 is unknown. **B.** Superposition of the yeast Aar2 (orange, PDB entry 3SBT; [47]) 3D structure and a homology model of human AAR2 derived from HHpred (marine), based on the yeast Aar2p structure [50]. The inset on the right shows a close-up view of the C-terminal domain, where serines 253 (yeast) and 284 (human) are displayed as sticks. Residues 154-170 and 318-356 of yeast Aar2 as well as residues 1-18, 170-201 and 359-384 of human AAR2 are not shown for clarity. **C.** Y2H interaction experiment with human AAR2 as bait and PRP8 as prey. Growth on selective media is abolished when AAR2 carries a S284E phospho-mimicry amino acid substitution. Mutation of second candidate phospho-site, AAR2(T356), does not influence the AAR2-PRP8 protein interaction. **D.** Interaction of AAR2 and PRP8 in human cells. HA-tagged PRP8 (aa1755-2335) was co-expressed in HEK293 cells with either wildtype AAR2, AAR2(S284A) or AAR2(S284E) mutant versions. Immunoprecipitation of PRP8 led to a co-precipitation of wild typeAAR2 and phospho-null AAR2(S284A). In contrast, the phospho-mimicry AAR2(S284E) version did not bind PRP8. **E.** Results of the phospho-Y2H screen to identify kinases modulating the interaction of AAR2 and PRP8. Growth on selective media (SD5), of strains carring AAR2 bait and PRP8 prey plasmids is reduced through co-expression of either SGK2 or CK2α1(CSNK2A1). Growth reduction is visible in a 1:10 dilution series on selective agar (SD5) in comparison to the control media (SD3). Ctrl: non-related kinases. **F.** Interaction of AAR2 and PRP8 in human cells in the presences of SGK and CK2α1 kinases. Experiment as in D. Co-expression of wild type SGK2 and CK2α1 reduces the amount of AAR2 co-immunoprecipitated with RPR8.

Here we investigate the phospho-dependent AAR2–PRPF8 interaction with the human proteins. The two proteins are conserved between yeast and human, however, it is not immediately evident from a standard sequence alignment whether the AAR2 phosphorylation sites including yeast serine 253 are conserved. The human AAR2–PRPF8 complex has been crystalized for high resolution structure determination [50]. We superimposed the two structures of the yeast and human Aar2/AAR2 protein and hypothesized that serine 284 in human may be functionally equivalent to serine 253 in yeast (**Figure 5B**). Notably, phosphorylation of the mammalian serine 284 is reported in PhosphoSitePlus [51]. Next, we recapitulated the interaction-disrupting effect of S253 phosphorylation observed in *S. cerevisiae* with the human orthologs using a Y2H approach (**Figure 5C**). Human PRPF8 bait (aa1755-2335) interacts with full length human AAR2 prey using our standard Y2H system [38,52]. While the S284A AAR2 mutant version has no effect, the S284E phospho-mimetic mutant in AAR2 abolished the Y2H growth readout (**Figure 5C**). In a control experiment, we assayed a second candidate phospho-site, T356A and T356E AAR2 mutant versions [47], also in combination with S284E, which confirmed that the S284 phospho-mimetic version of AAR2 was sufficient to abolish binding of PRPF8 in the human system (**Figure 5C**). When immunoprecipitating the human PRPF8 RNase H-like domain fragment from transfected HEK293 cells, both the wildtype AAR2 and the non-phosphorylatable S284A mutant versions showed comparable binding, while the S284E phospho-mimetic mutant completely abolished the interaction (**Figure 5D**).

In order to identify candidate human kinases that can mediate this phosphorylation-dependent PPI switch, we screened each of the 266 kinases for loss of the Y2H signal (**Figure 5E**). We transformed L40c **mat α**with the PRPF8-AAR2 bait and prey plasmids and mated this strain against the validated L40c **mata** kinase array. Each diploid strain expressed the two interacting proteins as well as one human kinase. After transfer on selective media we identified two candidate kinases SGK2 and CK2α1 (CSNK2A1), that reproducibly had a negative effect on Y2H reporter-dependent yeast growth (**Figure 5E**). Because Y2H utilizes a transcriptional reporter system, which has high sensitivity also to transient and less stable interactions [53], we do not expect that this setup will lead to complete loss of the signal. Rather, reduced, sub-stoichiometric phosphorylation of AAR2 may well account for the observed decrease in yeast growth of the SGK2 and CK2α1 strains.

To validate the kinase array phospho-Y2H results, AAR2 and PRPF8 were tested in a co-expression and immunoprecipitation assay in HEK293 cells. Co-expression of either SGK2 or CK2α1 reduced the amount of wild-type AAR2 co-immunoprecipitated with PRPF8 in comparison to the non-phosphorylatable mutant S284A (**Figure 5F**). Together, these experiments in mammalian cells support the results obtained with the kinase array when coupled to the phospho-Y2H assay. Active SKG2 and CK2α1 kinases substantially reduced the spliceosomal assembly complex interaction of AAR2 and PRPF8, and may thus be involved in triggering the interaction switch of PRPF8 from AAR2 to BRR2 in mammalian cells.

### Kinase-dependent interaction of ERα and 14-3-3 proteins

We next sought to test the utility of the kinase array in identification of kinases that promote protein-protein interactions. Estrogen receptor alpha (ERα), the hormone (β-estradiol) sensing nuclear receptor frequently overexpressed in breast cancer, is heavily post-translationally modified, including 26 phosphorylated residues [51]. Interestingly, De Vries-van Leeuwen et al. reported that ERα dimerization is inhibited through a phosphorylation-dependent interaction of the ERα monomer with 14-3-3 proteins. Mediated by the ERα C-terminal phospho-threonine 594, the interaction prevents the activation of ERα-dependent transcription (**Figure 6A)**, yet kinases which direct this inhibitory action remain elusive [54].

**Figure 6:**
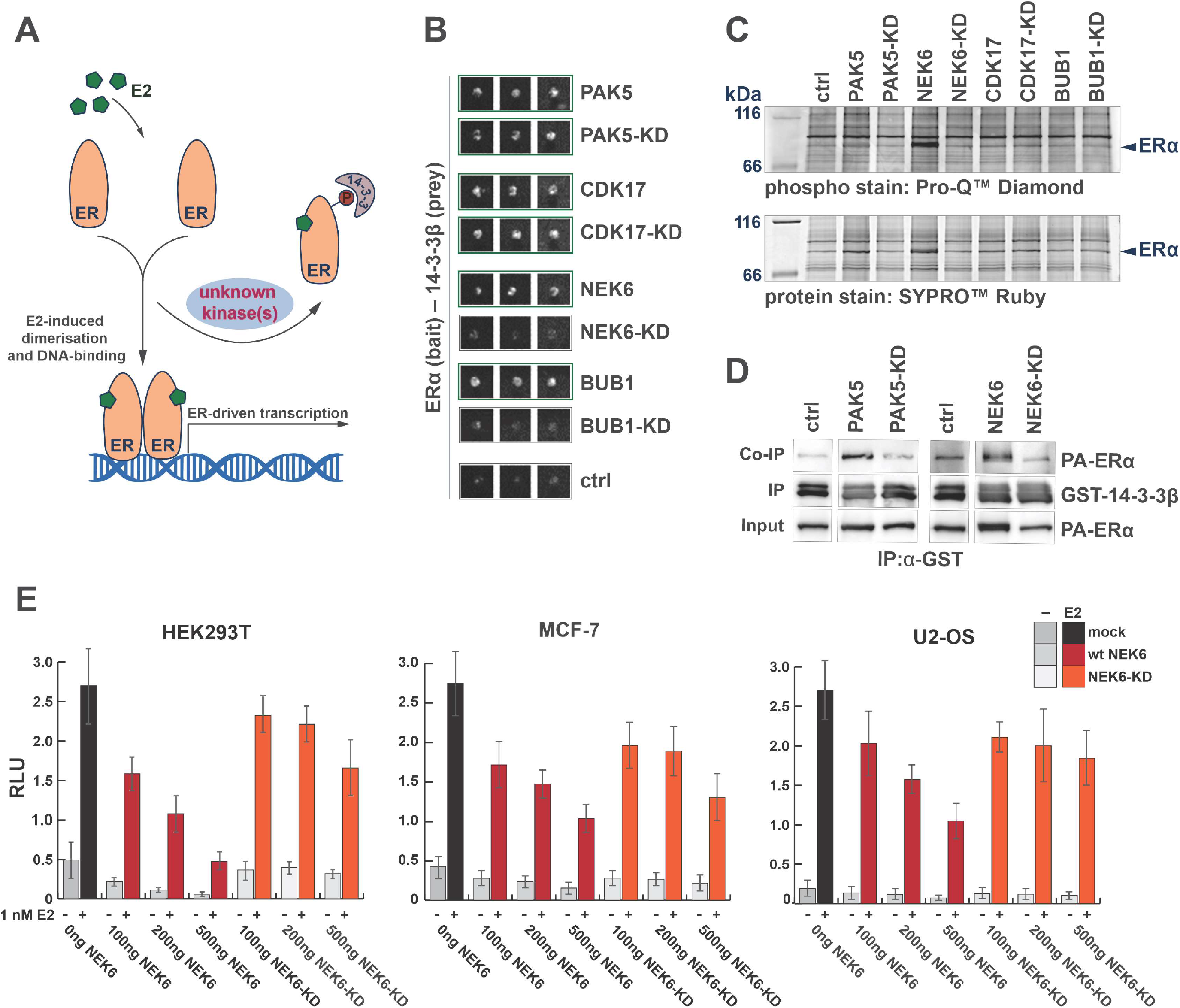
Identification of human S/T kinases modulating the ERα-14-3-3 phosphorylation-dependent interaction. **A.** Schematic of phospho-dependent interaction of the estrogen receptor ERα with 14-3-3β. Phosphorylation of ERα promotes its interaction with 14-3-3 proteins. This interaction removes ERα from the E2-stimulated ERα dimerization and as a consequence inhibits ERα chromatin interactions and gene expression. A protein kinase that can modulate the binding of ERα to 14-3-3 proteins is unknown. **B.** Results of the phospho-Y2H screen to identify kinases modulating the interaction of the ERα with 14-3-3β. The interaction of ERα (prey) with 14-3-3β (bait) is promoted by the co-expression of wildtype kinases of PAK5, CDK17, BUB1 and NEK6, for three biological replicas visible by the yeast growth on selective media in comparison to the control. The coexpression of the kinase-dead mutants of PAK5 and CDK17 also promoted the interaction of ERα and 14-3-3β. In contrast the kinase-dead mutant versions of BUB1(K821M) and NEK6(K74M&K75M) did not promote the Y2H interaction. **C.** Phosphorylation of ERα in mammalian cells. ERα was co-expressed with the indicated wildtype kinases or kinase-dead versions in HEK293 cells and immunoprecipitated. Phosphorylation status of ERα was analyzed with an in-gel phosphostain (Pro-Q^®^ Diamond phosphoprotein gel stain, Invitrogen), relative to the whole protein stain (SYPRO^®^Ruby protein gel stain, Invitrogen). **D.** Interaction of ERα and 14-3-3β in a GST pull-down assay. GST-tagged 14-3-3β was expressed in E. coli and immobilized on beads. PA-tagged ERα was co-expressed with either wildtype PAK5 or NEK6, the kinase-dead (KD) version or without a kinase in HEK293 cells. Immobilized 14-3-3β was used to pull PA-tagged ERα from the HEK lysates. Co-precipitation of ERα was successful in the presence of wild kinases, but not in their absence or with kinase-dead mutant versions. **E.** Effect of NEK6 co-expression on ERα gene activation. Relative luciferase activity of ERα in the presence and absence of 1 nM E2 (17β-estradiol) was measured. A dose-dependent reduction (ng plasmid transfected) in E2-Erα-dependent luciferase expression was observed in HEK293T, MCF-7 and US-OS cancer cell lines. This effect was not observed with the NEK6(K74M&K75M) kinase-dead version. Error bars indicate SD of triplicate transfections.

For phospho-Y2H analysis, we co-transformed L40c **matα** with ERα bait and 14-3-3 prey constructs for interaction mating with the human kinase yeast array. While ERα did not show interaction with 14-3-3β across multiple replicas in a classical Y2H experiment in the absence of a kinase (**Figure 6B**, Ctrl), the kinases BUB1, NEK6, CDK17, and PAK5 were able to reproducibly promote the interaction as indicated by the yeast growth on selective agar (**Figure 6B**). Parallel screening of NEK6(K74M&K75M), PAK5(K478M), BUB1(K821M), and CDK17(K221M) kinase-dead mutants suggested that phosphotransferase activity is required to promote the ERα – 14-3-3β interaction for the kinases NEK6 and BUB1. In contrast, kinase-dead mutants of PAK5 and CDK17 still promoted growth on Y2H selective medium (**Figure 6B**).

Up to 40% of the human kinome was predicted to bind to 14-3-3 proteins (including CDK17), and about 30 human protein kinases, including PAK5, have been shown to directly bind 14-3-3 proteins [55]. As PAK5 and CDK17 can potentially bind both ERα and 14-3-3, they may form a relatively stable ternary complex and therefore drive the positive Y2H signal in a phosphorylation-independent manner. To test this hypothesis, we performed a co-immunoprecipitation assay with a luciferase-based readout with 14-3-3β and wildtype as well as kinase-dead mutants of BUB1, NEK6, PAK5 and CDK17. The results indicated that the kinases PAK5 and CDK17 were able to bind to 14-3-3β while there no was binding detected for the wildtype or kinase-dead mutants of BUB1 or NEK6 (**Supplementary Figure 1** to 6B). Together with the phospho-Y2H results using kinase-dead mutants, these results are consistent with a model whereby PAK5 and CDK17 drive the Y2H interaction through formation of more stable ternary complexes with ERα and 14-3-3, while BUB1 and NEK6 promote ERα - 14-3-3 interaction mainly through phosphorylation.

To analyze whether the co-expression of the kinases BUB1, NEK6, PAK5 and CDK17 in HEK293 cells results in an increased phosphorylation of ERα, the phosphorylation status of ERα was determined with an in-gel phospho-stain. While no kinase activity of CDK17 and BUB1 could be detected on ERα after immunoprecipitation, the kinases NEK6 and PAK5 promoted the phosphorylation of ERα, in comparison to the corresponding kinase-dead mutants or the control sample without kinase overexpression (**Figure 6C**). A pulldown assay with bacterial-expressed GST-tagged 14-3-3β resulted in ERα binding if co-expressed with the wildtype kinases NEK6 or PAK5 in HEK293 cells when compared to the kinase-dead mutant co-expression or negative control experiments (**Figure 6D**). This result is consistent with the observed ERα-phosphorylation activity of NEK6 and PAK5 in HEK293 cells.

To summarize, NEK6 and BUB1 promote the association of 14-3-3β and ERα in Y2H in a kinase activity-dependent manner, and do not directly interact with 14-3-3β themselves. When co-expressed in mammalian cells, active forms of NEK6 and PAK5 increase the phosphorylation of ERα and its direct binding to 14-3-3β in GST pulldown assays. Together, these protein interaction data point to NEK6 as a prime candidate to regulate the association of ERα with 14-3-3β and thereby its transcriptional activity *in vivo*. To test this hypothesis, we transfected HEK293T cells with ERα and ERα-dependent luciferase reporter harboring three copies of Estrogen Response Element in front of a minimal promoter and a luciferase gene (3xERE-LUC, [56]), in presence of increasing amounts of wild-type or kinase-dead NEK6. The co-transfection of active NEK6 kinase inhibited estrogen-driven activation of the luciferase reporter in a dose-dependent manner (**Figure 6E**). The observed effect was dependent on the kinase activity of NEK6, as the attenuation of ERα transcriptional activity was negligible with the kinase-dead NEK6-KD version. This effect was reproduced in MCF7 cells, a breast cancer cell line and a standard model to study ERα transcriptional activity. Finally, also in U2-OS cells, the osteosarcoma cell line originally used by De Vries-van Leeuwen et al. to demonstrate the effects of 14-3-3 binding, active NEK6 kinase also reduced the ERα-dependent reporter gene transcription (**Figure 6E**). These results, obtained in three relevant mammalian cell lines, functionally support the results of our human kinase array screening approach, suggesting that NEK6 is a prime candidate kinase that can promote phosphorylation of ERα and its interaction with 14-3-3 proteins.

## Discussion

Kinase activity drives signaling processes and, when deregulated, human diseases. Kinases are the third largest group of human protein drug targets and kinase inhibitors represent the largest fraction of drugs with poly-pharmacological effects, that is they act through multiple targets [8]. However, beyond their primary sequence and domain architecture, a large fraction of human kinases are hardly characterized [9,10]. Systematic assaying multiple kinases activities in parallel is therefore fundamental to improve our understanding of cellular signaling, disease processes and drug action [9,57].

Experimental methods for assaying kinase activities include *in vitro* assays with recombinant proteins [57,58] or the application of kinase specific biosensors that work in cells [12–14]. For example, an enzyme assay panel employing more than 200 recombinant kinases was used with microfluidics capillary electrophoresis technology to screen for small molecule inhibitors [57]. Because results using recombinant kinases *in vitro* do not always recapitulate *in vivo* conditions, other approaches prove valuable to assay kinase activities. Taipale et al. [59] used the interaction of HSP90 / CDC37 chaperones with kinases as thermodynamic sensors of kinase activity or inhibitor binding in a cell culture co-IP approach. Also fluorescent biosensors examining conformational changes of kinase allow to temporally and spatially resolve kinase activities towards engineered targets in cells, tissues, and to some extent in whole organism [11,13,14]. Clearly, accessible tools to study active kinases under cellular conditions are a useful addition to current approaches.

This study presents information about the activity of 266 human protein kinases obtained from a yeast array, using growth as a readout [44]. While most kinases when expressed at low levels in yeast do not bestow a growth phenotype, we have increased sensitivity of the growth readout by employing a variation of growth conditions and by using four distinct yeast strains. Comparison to a set of non-active kinases and unrelated proteins indicated that although some phenotypes might be related to protein overexpression, this effect was not dominant. Moreover, some kinases may not be active in yeast because of the lack of regulatory proteins, or conversely, we may miss the activity of a set of kinases, where phosphorylation of yeast proteins simply is not detrimental to growth. For example, even though SGK2 and PAK5 were both grouped with the moderately active kinases, they could be identified from the array as candidates for modulating pS/T-PPIs because of the use of an alternative phospho-Y2H readout.

Examination of the yeast array yielded 150 highly active kinases. Our active kinase set is representative of the kinase families (**Figure 3**) and covers 35 members of the dark kinome [10], including CDK17, NEK6 and MARK3. The human kinase array can be leveraged for a variety of assays, provided the activity of a kinase can be coupled to a growth selection or other readouts. As proof of principle, we used the array to identify kinases that can modulate known pS/T-dependent protein interactions in a phospho-Y2H screen. While several approaches exist to define phospho-mediated PPIs, the kinases that modulate the PPI often remain elusive and so far, no systematic approach to identify them is available. First, we addressed the interaction of the spliceosomal protein PRPF8 and the snRNP assembly factor AAR2. Here we showed that phosphorylation of S284 of human AAR2 disrupts the interaction with PRPF8 (**Figure 5**), which in analogy to yeast is thought to constitute an important step in U5 snRNP and U4/U6-U5 tri-snRNP biogenesis [47,48]. The spliceosome is the large macromolecular protein-RNA complex that catalyzes pre-mRNA splicing. It assembles and disassembles on each pre-mRNA in a highly coordinated, stepwise manner governed by multi-protein-RNA interactions [52,60]. In yeast, cytoplasmic assembly of the U5 snRNP includes the assembly factor Arr2 in place of the spliceosomal helicase Brr2, which requires tight regulation [61]. There the binding of Arr2 to Prp8 blocks the Prp8 binding site of Brr2. As the Brr2-Prp8 interaction is blocked, Brr2 helicase activity may be shut off during this phase of U5 snRNP assembly. Nuclear phosphorylation of Aar2 disrupts its interaction with Prp8 and allows incorporation of Brr2 into U5 snRNP, and subsequently the U4/U6-U5 tri-snRNP through which Brr2’s helicase function is eventually delivered to the spliceosomal pre-catalytic B complex [48,49]. Subject to further scrutiny, a similar phosphorylation-dependent regulation of the AAR2-PRPF8 interaction is also thought to play a role in U5 snRNP or U4/U6-U5 tri-snRNP assembly in human. The kinase that could contribute in this regulation is presently unknown.

Screening the L40c yeast kinase array for a loss of the Y2H AAR2-PRPF8 bait-prey interaction revealed that the kinases SGK2 and CK2α1 are able to reduce the interaction (**Figure 5**). This result was confirmed by co-immunoprecipitation assays in HEK293 cells, where co-expression of wild type SGK2 or CK2α1 reduced the amount of AAR2 co-precipitated with PRP8. The AGC kinase group member SGK2 is not characterized very well. SGK kinases, just like AKT/PKB kinases are activated by PDK1 through phosphorylation of conserved residues [62]. SGK2 is known to be involved in the phosphorylation of diverse set of cellular channels and receptors [63] including V-ATPase proton pump [64]. The other candidate CK2α1, however, is a constitutively active kinase primarily located in the nucleus and thus could support disruption of the AAR2-PRPF8 interaction in the nucleus during late stages of U5 or U4/U6-U5 assembly. CK2α1 has been reported to phosphorylate a vast number of substrates and to regulate numerous cellular processes [65–67], including cell cycle progression, apoptosis and transcription, as well as a viral infection. It is involved in other spliceosomal processes such as the phosphorylation of Prp3 [68,69], the exon junction complex member / splicing activator RNPS1 [70] and is associated with the U4/U6/U5 tri-snRNP complex [67].

Estrogen receptor alpha (ERα / ERS1) is one of the most highly studied hormone sensing nuclear receptors. Its dimerization is essential for its ligand-dependent transcriptional activity. Blocking ERα function is the major route to treat luminal (ER+) breast cancer. ERα is phosphorylated at various sites, and potential functions and protein kinases for some phosphorylation sites have been characterized [71]. A phosphorylation-dependent interaction between ERα and the phospho-binding adaptors 14-3-3, which reduces the transcriptional activity and the proliferative effect of ERα in cells has been reported. The authors specifically demonstrated that phosphorylation of ERα is required for the binding of 14-3-3 protein and proposed that stabilization of this interaction with its effect on ERα transcriptional activity may be of therapeutic value [54].

We identified four candidate kinases, BUB1, NEK6, PAK5 and CDK17, with the ability to promote the interaction of ERα with 14-3-3β in a phospho-Y2H matrix approach from screening the kinase L40c array (**Figure 6**). In this assay, kinase activity of BUB1 and NEK6 was required to promote the interaction, whereas kinase inactive versions of PAK5 and CDK17 were also sufficient for a positive Y2H growth readout. The latter results are explained by the ability of the two kinases to stably bind to 14-3-3 proteins. This binding is reported in the literature [55] and was confirmed in luciferase-based co-immunoprecipitation experiments. However, we detected increased phosphorylation of ERα phosphorylation in HEK293 cells upon co-expression of wild type NEK6 and PAK5 kinase. Expression of these two kinases also led to successful co-immunoprecipitation of ERα with 14-3-3β. In order to demonstrate that NEK6 phosphorylation-dependent increase of ERα–14-3-3 interaction can have functional consequences in cells, we performed ERα-dependent transcription activation assays. In all three cell lines tested, HEK293, the breast cancer cell line model MCF-7 and U2-OS cells, overexpression of the active NEK6 kinase reduced transcriptional activity in a dose-dependent manner, an effect not observed with the kinase inactive NEK6 variant.

Among the four candidate kinases with the potential to modulate ERα activity via a phosphorylation-dependent interaction with 14-3-3 proteins we have identified, BUB1 is a fairly well characterized kinase involved in spindle checkpoint control [72]. In contrast, PAK5, CDK17 and NEK6 belong to the group of understudied, less well characterized kinases. PAK5 (formerly PAK7) is an actin and microtubule associated p21^cdc42/rac1^ - activated kinase family member [73], which has been shown to promote proliferation, invasion and migration of various cancer cell models [74]. CDK17 (also PCTAIRE2) is an uncharacterized member of the cyclin-dependent kinase family [75]. Taken together, our data point to NEK6, a cell cycle kinase involved in progression through mitosis [76,77], with only few substrates known so far [78], as the strongest candidate as ERα effector.

## Conclusion

Here, we built on the observation that heterologous expression of human kinases in yeast can cause a growth defect attributed to the enzymatic phosphotransferase activity of these proteins [29,30,44]. We have demonstrated that a large fraction of human kinases (56%) is active in our yeast array. For a subset of kinases, we have also demonstrated the kinase activity directly using kinase-dead mutant version as comparisons, or through monitoring the extent of yeast protein phosphorylation with phospho-specific antibodies. Mechanism that explain why aberrant phosphorylation of yeast proteins impar yeast growth are largely elusive and may be different from kinase to kinase. Whether or not kinase activity can be reported using the yeast array depends on growth conditions, the yeast strain and the readout used. Readouts where kinase activity is coupled to growth inhibitory effects, either directly or via a reporter system, can be used to study kinase perturbation [23] and inhibition [18,30,79,80]. Under conditions of low kinase expression levels, that do not impair growth, different setups can be utilized to pick up kinase activities, for example mass spectrometry-based substrate identification [5] or screens for pY-dependent protein interactions [19]. Here, we extended the phospho-Y2H approach to S/T kinases for the first time, utilizing the array to identify kinases that can modulate known pS/T-dependent protein interactions. Knowledge of potential kinases that regulate phospho-dependent protein interactions, many of which are prime drug targets, can lead to novel pharmacological intervention strategies. Therefore, the human kinase array approach is a versatile methodological addition, allowing to perform comprehensive studies of human kinases and their effects.

## Material and Methods

### Antibodies

The following antibodies were used: mAB anti-phospho-CDK substrate (pTPXK, rabbit, Cell Signalling, 14371), mAB anti-phospho-CK2 substrate MultiMab™ ((pS/pT)DXE, rabbit, Cell Signalling, 8738), mAB anti-phospho- MAPK/CDK substrate (PXS*P or SPXR/K, rabbit, Cell Signalling, 2325), mAB anti-phospho-PKA substrate (RRXS*/T*, rabbit, Cell Signalling, 9624), pAB anti-phospho-(Ser)-PKC substrate ((R/K)X(S*)(Hyd)(R/K), rabbit, Cell Signalling, 2261), pAB anti-GST (goat, GEHealthcare, 27457701V), mAB anti-HA (mouse, Hiss Diagnostics GmbH, AB-10110), mAB anti-phospho-tyrosine clone 4G10 (mouse, Millipore, 05-321), secondary mAB anti-goat HRP (rabbit, Invitrogen, 611620), secondary mAB anti-mouse (sheep, GE Healthcare, LNA931V), secondary mAB anti-rabbit (donkey, GE Healthcare, LNA934V);

### Yeast strains

BY4742 (MATα) MATα his3Δ1 leu2Δ0 lys2Δ0 ura3Δ0, [35,36]; L40c (MATa) his3Δ200 trp1-901 leu2-3,112 LYS2::(lexAop)4-HIS3 ura3::(lexAop)8-lacZ ADE2::(lexAop)8-URA3 GAL4 gal80 can1 cyh2 [37,38]; W303 (MATa/MATα) leu2-3,112 trp1-1 can1-100 ura3-1 ade2-1 his3-11,15 [39]; Y258 (MATa) pep4-3 his4-580 ura3-52 leu2-3 [40];

### Growth phenotype screen

Supplementary Table 1 lists all conditions successfully tested including concentrations of additives. The four yeast strains were transformed with 1,020 clones (pASZ-C-DM or pASZ-CN-DM) representing kinases and protein set. For the growth phenotype screen, the kinase yeast array was transferred in 384 format using a gridding robot and grown on agar plates with minimal medium with 2% glucose and 20 μM CuSO_4_ at 30°C unless otherwise stated (6-8 days). Yeast growth strength was visually evaluated in comparison to the overall growth of the yeast strain in the one condition and recorded using a 0-3 scale. For each of the four strains, plotting the number of conditions over the number of kinases with growth phenotype gave a bimodal distribution separating conditions with relatively fewer specific kinase signals from conditions that have a general diminishing effect on yeast growth. According to this distribution, conditions which reduced growth of more than 175 colonies were excluded from further analysis.

### Phospho-Y2H Screen

The general Y2H set-up has been described in detail previously [38]. For the phospho-Y2H screen, the L40cα Y2H strain was co-transformed with the plasmids expressing the bait constructs (pBTM116-D9 or pBTMcC24-DM) and the prey constructs (pCBDU-JW or pACT4-DM). The independently transformed yeast colonies were mated with the L40c kinase array strains on YPD agar (30°C, 2 days). Active kinases modulating the phosphorylation-dependent PPI were identified by growth on selective media (SD5) supplemented with 20 μM or 100 μM CuSO_4_ for the induction of kinase expression (30°C, 3-5 days).

### Yeast growth curves

To measure yeast growth in liquid culture, triplicates of the W303 strain expressing a single human protein kinase were created on agar, transferred to liquid media in microtiter plate format. For the growth measurement, a 96-well MTP was incubated for 1h at 30°C with orbital shaking, the yeast was then diluted with selective media (SD5, 500 μM CuSO_4_) with or without 3.2% (v/v) DMSO to a starting OD_595_ of 0.10-0.15. Then yeast growth was recorded in the MTP for 24 h by taking the OD every 10 min and shaking every hour for 5 min.

### Protein interaction assays in mammalian cells

HEK293T cells were cultured in Dulbecco′s modified eagle medium (DMEM) with 10% fetal bovine serum (FBS) in a humidified 5% CO 2 atmosphere at 37°C. For the immunoprecipitation and pull-down experiments ~6 ×10^7^ HEK293T were cells plated in a 10 cm cell culture dish and incubated for 24 prior to transfection. A total of 4 μg DNA of the different constructs were used for transfection with Lipofectamine 2000.

### GST pull-down experiments form HEK293T cells (AAR2-PRPF8 interaction)

GST-tagged AAR2 (pcDNA5/FRT/TO-nGST) and the HA-Strep-tagged PRPF8 (pcDNA5/FRT/TO/SH/GW) constructs were co-transfected with PA-tagged kinase in HEK293T cells. HA-Strep-tagged PRPF8 protein was pulled down with Strep-Tactin Sepharose and the amount of bound AAR2 protein was determined by western blotting. Cells were harvested 24 h after protein expression induction lysed in 0.5 ml LyseH-buffer (50 mM HEPES, 500 mM NaCl, 10% Glycerol, 0.5 mM EDTA, 2 mM MgCl_2_, 1% NP-40, 12.1 mM sodium deoxycholate, 1x Phosphatase inhibitor III, 1x Protease inhibitor, 0.1 U/μl Benzonase), cleared and incubated with human Strep-Tactin sepharose beads for 1 h on ice. The beads were washed four times with ice-cold WashH buffer (50 mM HEPES, 500 mM NaCl, 10% Glycerol, 0.5 mM EDTA, 2 mM MgCl_2_, 1% NP-40, 12.1 mM sodium deoxycholate) and treated two times with 30 μl 2x SDS loading buffer for SDS-PAGE western blot analyses.

### Immunoprecipitation of ERα

To analyze ERα phosphorylation in mammalian cells, PA-tagged ERα (pcDNA3.1PA-D57) was co-expressed with a GST-tagged kinase (pcDNA5/FRT/TO-nGST) in HEK293T cells, immunoprecipitated and the phosphorylation status of ERα was analyzed with Pro-Q phospho-stain (Thermo Scientific) of SDS-PAGE. Transfected HEK293T cells were harvested 24 h after protein expression induction and lysed in 0.5 ml LyseH-buffer for 30 min on ice. The cleared cell lysate was transferred onto pre-washed 30 μl human IgG agarose beads and incubated 1 h on ice. The beads were washed four times with ice-cold WashH buffer and proteins were eluted with 30 μl 2x SDS loading buffer. After SDS gel electrophoresis, the gels were stained with Pro-Q Diamond phospho-stain and subsequent SYPRO Ruby whole protein stain and scanned with a fluorescence scanner according to the manufacture’s instructions.

### GST pull-down experiments form HEK293T cells (ERα-14-3-3 interaction)

GST-tagged 14-3-3 constructs (pDESTcoG) were expressed in E. coli, then loaded on GST-coated beads and incubated with HEK293T cell lysate containing co-expressed PA-tagged ERα and GST-tagged kinases. The amount coprecipitated ERα was detected by western blotting. GST-tagged 14-3-3 was expressed in 12 ml E. coli culture (TB-medium, 100 μg/ml ampicillin, 30 μg/ml chloramphenicol) for about 20 h at 37°C and 150 rpm shaking. E. coli cells were lysed in 1.85 ml LyseB buffer (50 mM HEPES, 5400 mM NaCl, 5% Glycerol, 1 mM EDTA, 0.5% Brij58, 1mg/ml Lysozyme, 2mM DTT). 0.1U/μl Benzonase was added in 50 mM HEPES pH 8, 2mM MgCl2 and incubated for 30 min at 4°C. 9 μl supernatant with expressed 14-3-3β were used for the pulldown experiment with 5 μl human Glutathione magnetic beads pre-incubated with 100μl LyseH buffer each. The HEK293T cell lysates were prepared 24 h after protein expression induction in 0.5 ml LyseH-buffer mixed with the GST-fusion beads with the bound 14-3-3 proteins and incubated on ice for 1h. The beads were washed six times with 0.5 ml ice-cold WashH buffer and samples were eluted from the beads with 30 μl 2x SDS loading buffer. Samples were subject to SDS gel electrophoresis and subsequent western blotting.

### Co-Immunoprecipitation with luciferase readout

Luciferase based co-immunoprecipitation experiments were performed in HEK293T cells in 96 well format using 150 ng of plasmid DNA in total (firefly-V5 fusion 14-3-3β construct in pcDNA3.1V5Fire-DM and kinase protein A fusion constructs in pcDNA3.1PA-D57) as described previously [52,81].

### Transient transfection luciferase transcription assays

The cells were seeded out in phenol red – free DMEM supplemented with 10% charcoal stripped FBS on 96-well plates. 24 hours after seeding the cells were transfected with indicated plasmids using PEI methods with PEI to DNA ratio of 5:1 w/w or 2.5:1 for HEK293T and MCF7 or U2-OS, respectively. Six hours after transfection, the cells were stimulated with 1 nM E2 (17β-estradiol). 24h after transfection, the cells were washed and lysed in Luciferase Cell Culture Lysis Reagent (Promega). The luciferase activity was quantified using Luciferase Assay System (Promega) and normalize to the protein content.

### Data processing and visualization

Data processing, statistical analyses and visualization was performed in R, kinome tree graphics were produced using CORAL at http://phanstiel-lab.med.unc.edu/CORAL/.

## Author Contributions

Conceptualization: US

Formal Analysis: SJ, NK; JW, US

Funding acquisition: US, MW

Investigation: SJ, NK, NB, GW

Methodology: SJ, JW, US

Resources: US, MW

Supervision: US, MW

Validation: SJ, NB, NK

Visualization: NK, SJ, JW, US, GW

Writing – original draft: US

Writing – review & editing: NK and all authors

## Acknowledgments

The work was supported by the Max Planck Society and the University of Graz.

---

**Supplementary Table 1: Conditions**

Information about 73 different conditions (33 unique treatments) which were successfully evaluated for at least one of the four strains. A total 243 condition – strain pairs were considered.

**Supplementary Table 2: Kinases**

Information about the 266 human kinases assayed for activity in yeast. Activity ranks are based on binning the number of conditions where growth reduction was observed in each strain (0-4) followed by clustering (Figure 1). Annotations shown in Figure 3 are listed.

**Supplementary Figure 1:**
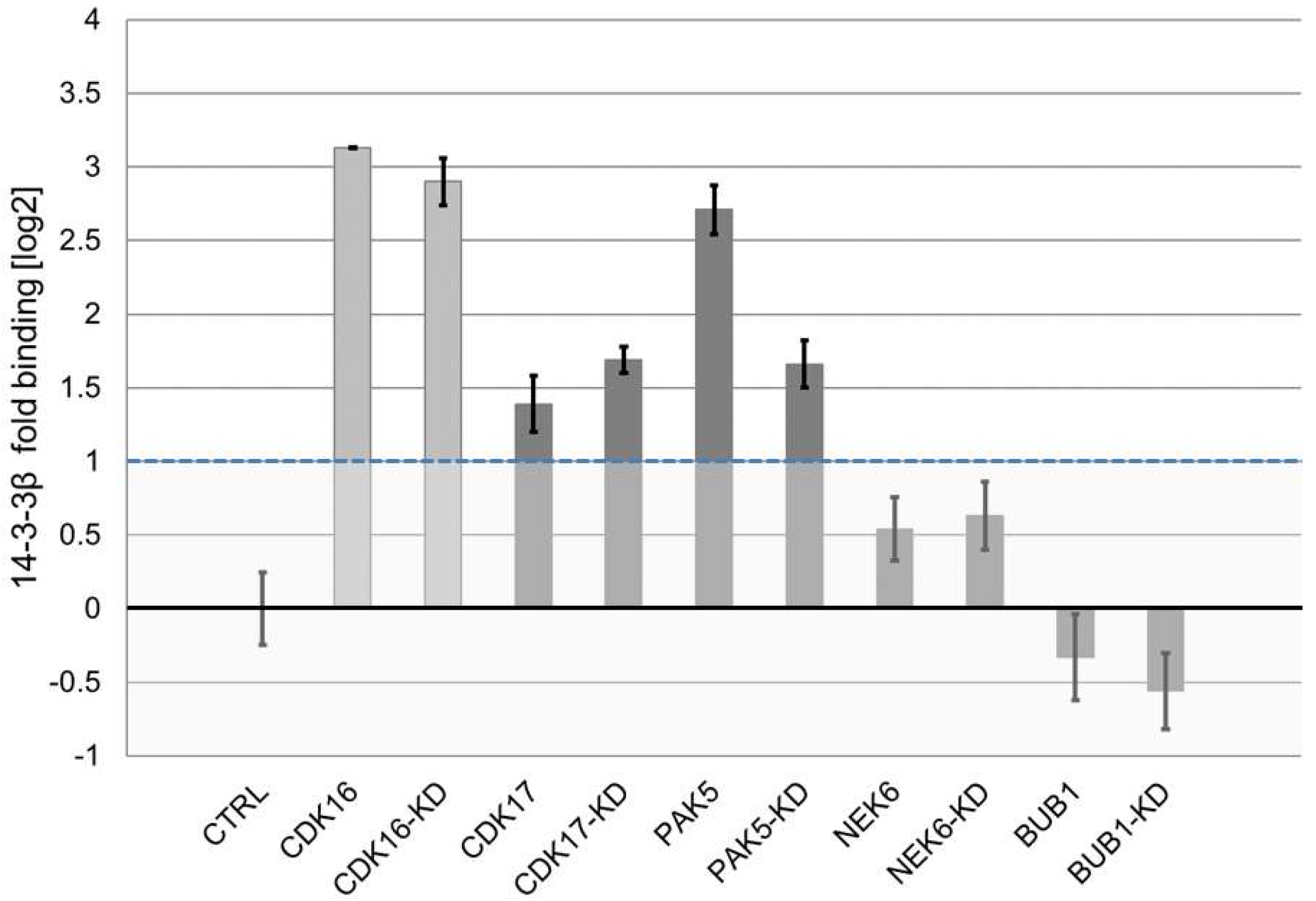
Luciferase based Co-immunoprecipitation experiments. Results of the LUMIER assay with 14-3-3β (tagged with firefly luciferase) and the indicated kinases (protein A tagged) identified in the Y2H-Screen. CDK16 is a known interaction partner of 14-3-3β and served as positive control (light grey). Error bars indicate SD of triplicate transfections.

